# Matrisome AnalyzeR: A suite of tools to annotate and quantify ECM molecules in big datasets across organisms

**DOI:** 10.1101/2023.04.18.537378

**Authors:** Petar B. Petrov, James M. Considine, Valerio Izzi, Alexandra Naba

**Affiliations:** Infotech Institute, University of Oulu, FI-90014 Oulu, Finland; Department of Physiology and Biophysics, University of Illinois Chicago, Chicago, IL 60612, USA; Faculty of Biochemistry and Molecular Medicine & Faculty of Medicine, BioIM Unit, University of Oulu, FI-90014 Oulu, Finland; Foundation for the Finnish Cancer Institute, Tukholmankatu 8, Helsinki, Finland; University of Illinois Cancer Center, Chicago, IL 60612, USA

**Keywords:** Extracellular Matrix, Bioinformatics, Omics, Data Annotation, Model Organisms

## Abstract

The extracellular matrix (ECM) is a complex meshwork of proteins that forms the scaffold of all tissues in multicellular organisms. It plays critical roles in all aspects of life: from orchestrating cell migration during development, to supporting tissue repair. It also plays critical roles in the etiology or progression of diseases. To study this compartment, we defined the compendium of all genes encoding ECM and ECM-associated proteins for multiple organisms. We termed this compendium the “matrisome” and further classified matrisome components into different structural or functional categories. This nomenclature is now largely adopted by the research community to annotate -omics datasets and has contributed to advance both fundamental and translational ECM research. Here, we report the development of Matrisome AnalyzeR, a suite of tools including a web-based application (https://sites.google.com/uic.edu/matrisome/tools/matrisome-analyzer) and an R package (https://github.com/Matrisome/MatrisomeAnalyzeR). The web application can be used by anyone interested in annotating, classifying, and tabulating matrisome molecules in large datasets without requiring programming knowledge. The companion R package is available to more experienced users, interested in processing larger datasets or in additional data visualization options.

**SUMMARY STATEMENT:** Matrisome AnalyzeR is a suite of tools, including a web-based app and an R package, designed to facilitate the annotation and quantification of extracellular matrix components in big datasets.

## INTRODUCTION

The extracellular matrix (ECM) is a complex meshwork of proteins that forms the scaffold of all multicellular organisms (Hynes and Naba, 2012). It plays critical roles in all aspects of life: from orchestrating cell migration and differentiation during development (Dzamba and DeSimone, 2018; Walma and Yamada, 2020), to supporting tissue growth and repair. It also plays critical roles in the etiology or progression of diseases (Theocharis et al., 2019).

Omic technologies (*e*.*g*., transcriptomics, proteomics, glycomics) have emerged as powerful approaches to profile at large-scale, and often in an unbiased manner, the biomolecular landscape of cell and tissue states. However, to extract meaningful information and generate novel hypotheses, we need to develop comprehensive annotations and analytical methods to mine these complex inputs. Hence, to study the ECM using -omic technologies, we first needed a compendium of all potential ECM components. Using *de-novo* sequence analysis (Gebauer and Naba, 2020; Naba et al., 2012b; Naba et al., 2016), we have predicted the “matrisome” of multiple organisms, including human (Naba et al., 2012a), mouse (Naba et al., 2012a), zebrafish (Nauroy et al., 2018), fruit fly (Davis et al., 2019), and nematode (Teuscher et al., 2019). We further classified matrisome genes into: 1) the “core matrisome”, *i*.*e*., the genes encoding structural components of the ECM including ECM glycoproteins, collagens, and proteoglycans, and 2) “matrisome-associated”, *i*.*e*., the genes encoding non-structural components of the ECM that either share structural similarities with core matrisome components (we termed these “ECM-affiliated proteins”), or are capable to modulate the structure (“ECM regulators”) or signaling (“secreted factors”) functions of the ECM proper (**Table 1**).

The matrisome lists have been deployed via different platforms to support data analysis including the Molecular Signature Database (Subramanian et al., 2005), the Zebrafish Information Network (Bradford et al., 2022), or FlyBase, the database of *Drosophila* Genes and Genomes (Gramates et al., 2022). Used to annotate transcriptomic datasets, these matrisome lists have contributed, for example, to help identify the diverse cell populations expressing ECM genes in health an diseases (Bergmeier et al., 2018; Etich et al., 2019; Nauroy et al., 2017; Pietilä et al., 2021; Wietecha et al., 2020) or to identify networks of ECM genes characteristic of disease stages or of prognostic value (Izzi et al., 2018). Used to annotate proteomic datasets, these lists have enabled the definition of the ECM composition of tissues and organs across the pathophysiological spectrum (Naba, 2023; Randles et al., 2017; Shao et al., 2023).

To facilitate the use of the matrisome classification, we previously developed a web application capable of handling human and murine proteomic datasets (Naba et al., 2017). The previous iteration required users to extensively format their input datasets to be amenable, which hindered its diffusion to recently-growing methodologies such as single-cell RNAseq (sc-RNAseq). Here, we report the development of Matrisome AnalyzeR, an augmented suite of versatile tools that includes a web-based Shiny application (https://sites.google.com/uic.edu/matrisome/tools/matrisome-analyzer) and a companion R package (https://github.com/Matrisome/MatrisomeAnalyzeR). The new intuitive web-based application can be used by anyone to obtain the annotation, classification, and tabulation of matrisome molecules from their datasets. In the Matrisome AnalyzeR app, results appear on screen in seconds and change dynamically in response to user actions, through a user-friendly, point-and-click interface requiring no programming knowledge. The companion Matrisome AnalyzeR package is available to more advanced users interested in processing larger files (>30MB) and provides additional data visualization options and possibilities for integration with more complex pipelines. In their current versions, the Matrisome AnalyzeR web application and R package are capable of processing data from the following organisms: *Homo sapiens* (human), *Mus musculus* (mouse), *Danio rerio* (zebrafish), *Drosophila melanogaster* (fruit fly), and *Caenorhabditis elegans* (roundworm).

## RESULTS

### The web-based Matrisome AnalyzeR Shiny application

#### Data input

**Fig. 1A** illustrates the data input process. Users can upload tab- or comma-separated (.tsv, .txt, .csv) files with column headers, not exceeding 30MB. If a data file exceeds this limit, we recommend using the Matrisome AnalyzeR package *(see below*). A test file containing proteomic data from three technical replicates on ECM samples from human Fallopian tubes (Renner et al., 2022) is provided as **Supplementary Table S1** and will be used as an example here. The test file is also available for download via the web-based Shiny application and included with the R package.

**Figure 1.**
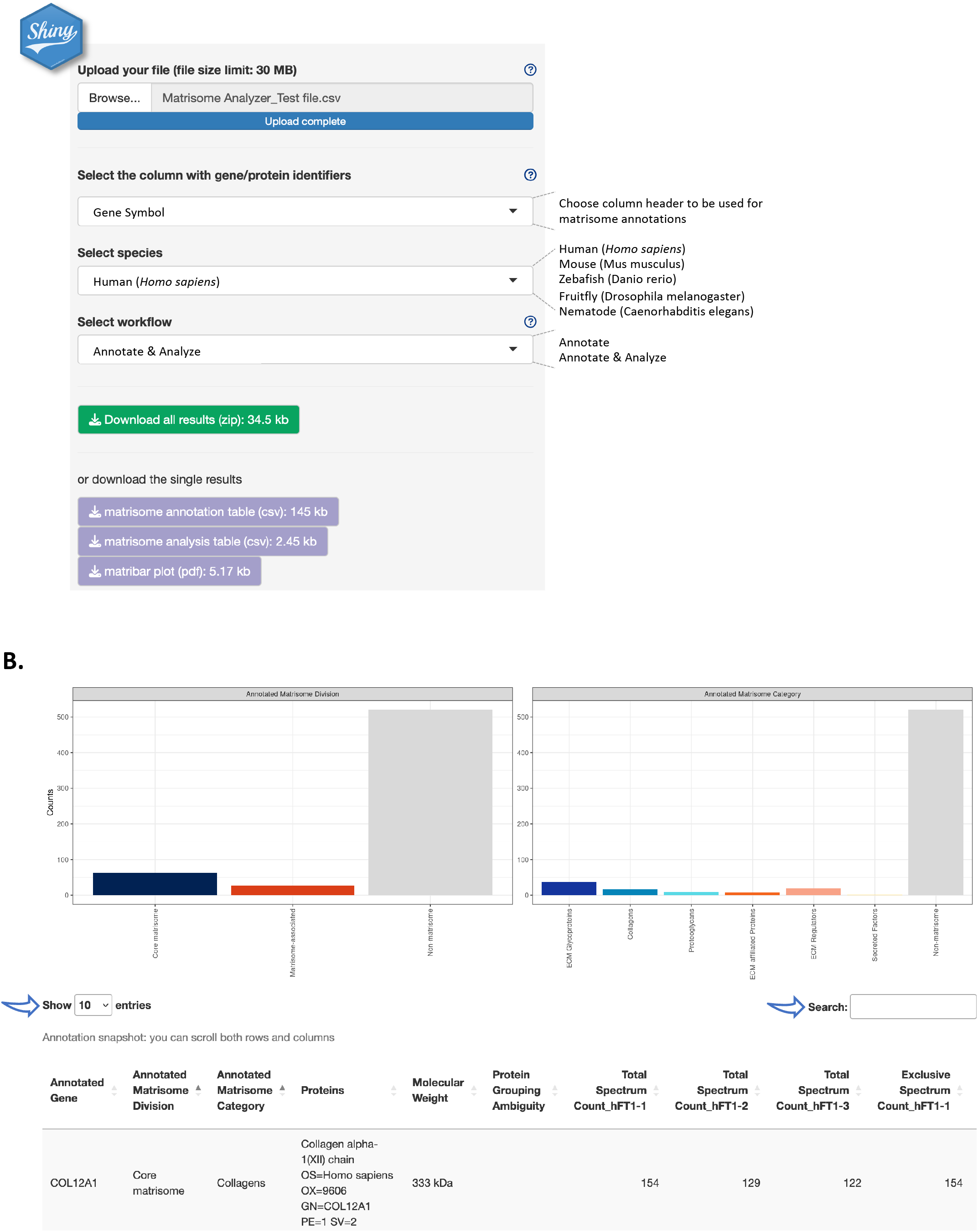
The web-based Matrisome AnalyzeR Shiny application interface. **A**. Home page of the Matrisome AnalyzeR web application (https://sites.google.com/uic.edu/matrisome/tools/matrisome-analyzer) displaying input parameter options and output files. **B**. Running the “annotate + analyze workflow” using Supplementary File S1 as input, returns bar graphs (or “`matribars`”) representing the total numbers of matrisome molecules (here, proteins) classified according to matrisome divisions (left panel) and matrisome categories (right panel) across the entire dataset and a searchable and customizable table (arrows).

Upon file upload, Matrisome AnalyzeR will automatically recognize number format, but we encourage using dots, and not commas, for decimals and avoiding formatting thousands. Matrisome AnalyzeR will also automatically populate the first box with column headers. Users will be asked to select, from the next two drop-down menus, the column containing the molecule identifiers to be used for the annotation and the species of interest. The tool is currently designed to accept gene symbols, NCBI gene (formerly Entrez Gene), and UniProt IDs (The UniProt Consortium, 2023) for all species. Additionally, Matrisome AnalyzeR accepts Ensembl Gene IDs for human and murine datasets, ZFIN IDs for zebrafish datasets, FlyBase ID for *Drosophila* datasets, and WormBase and Common Gene Name for *C. elegans* datasets. In the eventuality that no identifiers map to the application’s database, an error message will prompt users to review input choices. After input selection, users will then select the workflow to process their data. Help buttons have been implemented to further facilitate data input (**Fig. 1A**).

#### Data annotation

The “Annotate” workflow annotates the input file with matrisome divisions (*i*.*e*., Core matrisome, Matrisome-associated, or Non-matrisome) and categories (*i*.*e*., ECM glycoproteins, Collagens, Proteoglycans, ECM-affiliated, ECM regulators, Secreted Factors, or Non-matrisome). The output provides a .csv file that corresponds to the original input file, where column A lists the identifiers used for the annotation in alphabetical order, column B, the “Annotated Matrisome Division”, and column C, the “Annotated Matrisome Category” (**Supplementary Table S2**). The output table is also visible and browsable on the main page upon completion of the “Annotate” workflow (**Fig. 1B**). Users can customize the number of entries displayed in the main window and can search the table using the search box (**Fig. 1B**). In addition, the output includes a .pdf file with bar graphs (or “`matribars`”) representing the total numbers of matrisome molecules (*e*.*g*., genes, proteins) classified according to matrisome divisions and categories across the entire dataset (**Supplementary File S1**). These bar graphs are also displayed on the main page upon completion of the “Annotate” workflow (**Fig. 1B**). Note that the output can change dynamically in response to user actions, through the user-friendly point-and-click interface.

#### Data analysis

The “Annotate + Analyze” workflow does the above and then tabulates the content of each numerical column in the input by matrisome divisions and categories. Here, the output is a .csv file where each row corresponds to a matrisome classification, where column A lists “Matrisome Annotations”, and where each subsequent columns report the tabulation of the numerical data according to these annotations (**Supplementary Table S3**). This workflow allows users to evaluate, at a glance, the relative ECM content of each of their samples (*e*.*g*., number of reads if inputting RNA-seq data or number of peptides or spectra if inputting proteomic data, as shown in the example provided in **Supplementary Table S1**). Users can then input these data in other statistical analysis software or data visualization software to pursue their analysis.

We caution users that not all tabulations might be relevant: for example, the test file provided contains representative proteomic data, listing in addition to quantitative metrics (*e*.*g*., Total Spectrum Count or Exclusive Spectrum Count), the molecular weight of each protein or protein identification probabilities that are numerical values and, thus, are tabulated by Matrisome AnalyzeR.

Importantly, the Matrisome AnalyzeR application implements a strict session-specific data policy: data uploaded by users are neither stored in our server, nor can the data leak through sessions. User data are purged upon user disconnection or at session timeout.

### The Matrisome AnalyzeR package

For users familiar with R programming and wishing to analyze larger datasets (>30MB, the limit imposed on data upload to the web application) or interested in additional data visualization options, as well as, in the possibility of integrating matrisome annotation and analysis with other existing analysis pipelines, we have developed the “Matrisome AnalyzeR” R package, available at: https://github.com/Matrisome/MatrisomeAnalyzeR.

The Matrisome AnalyzeR GitHub repository includes all the functions required to run the data processing workflow from the annotations to the tabulations as described above for the web application. Additional data visualization options are available such as donut charts (“`matrirings`”), polar bar charts (“`matristars`”), or alluvial charts (“`matriflows`”). **Fig. 2A** shows examples of these visualization options for the data provided as test file (**Supplementary Table S1**).

**Figure 2.**
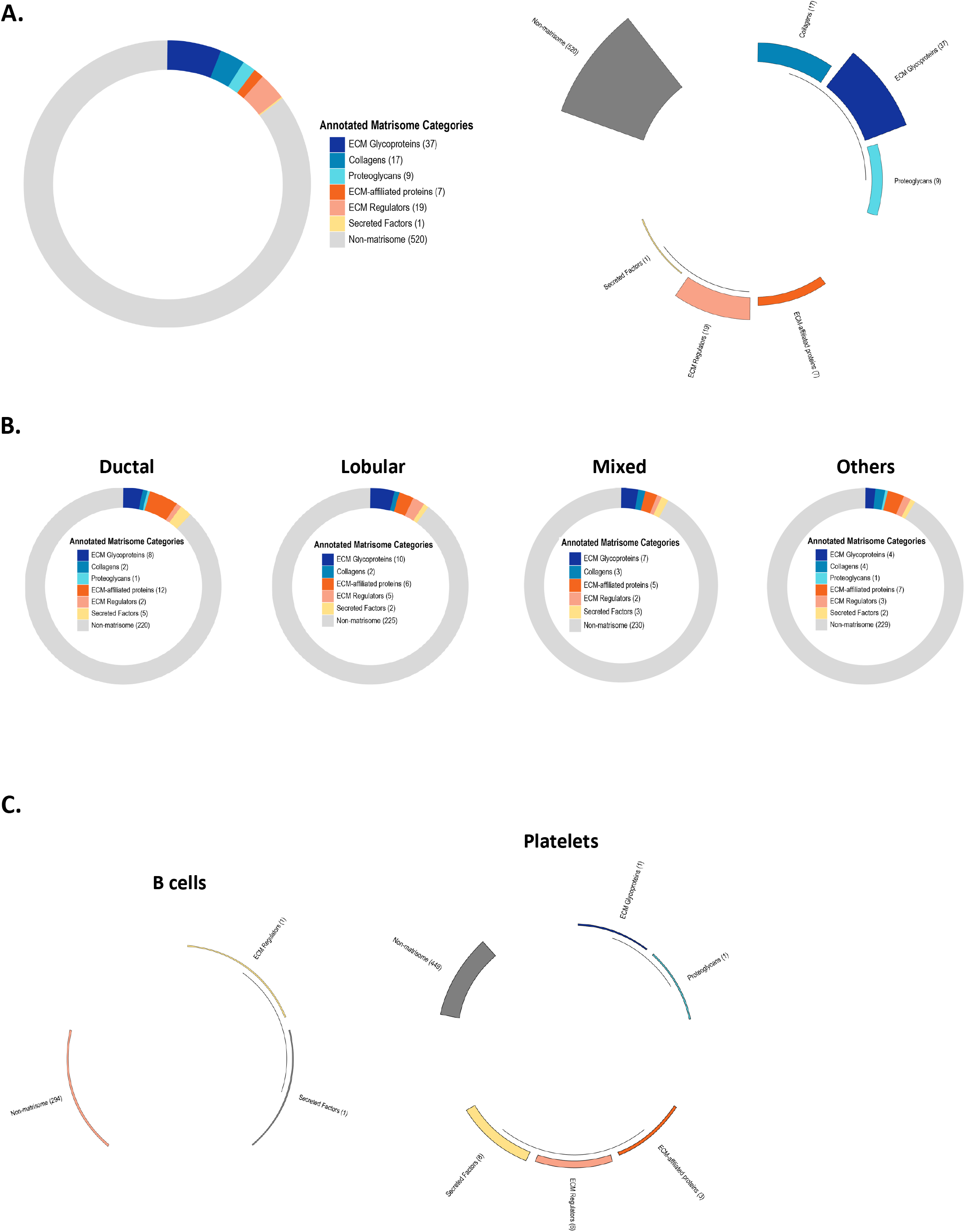
Additional data visualization options using the Matrisome AnalyzeR package. **A**. Upon running the “`matriannotate`” function on Supplementary Table S1 as the input, users can obtain additional output files representing the data as donut chart (“`matriring`”; left panel) or polar bar chart (“`matristar`”; right panel). **B**. Donut charts (or “`matriring`”), obtained using the Matrisome AnalyzeR package to analyze whole exome sequencing data on four classes of breast cancer retrieved from the cBio portal, represent the 250 genes presenting the highest mutation frequencies in each breast cancer subtypes and their classification into matrisome categories. **C**. Polar bar charts (“`matristars`”), obtained using the Matrisome AnalyzeR package to analyze single-cell RNA-seq data from 2,700 single peripheral blood mononuclear cells, represent the average expression level of matrisome and non-matrisome gene categories for each single cell cluster such as B cells (left panel) and platelets (right panel). Differences in gene expression levels between categories and across cell clusters are visualized through the length of each segment and height of each segment’s bar.

The GitHub repository also features additional case studies to demonstrate the breadth of Matrisome AnalyzeR. In a second example, we show how the Matrisome AnalyzeR package can be applied to the analysis of whole exome sequencing data obtained from the cBioPortal (Cerami et al., 2012; Gao et al., 2013) to identify matrisome genes presenting a high mutation frequency across four different breast cancer subtypes. Processing of the dataset using Matrisome AnalyzeR results in donut charts representing, in our example, the 250 genes presenting the highest mutation frequencies in each breast cancer subtypes and their classification into matrisome categories (**Fig. 2B**). By selecting the donut chart representation (or “`matriring`”), users can easily visualize the contribution of matrisome genes to the query and for example identify a switch from the predominantly ECM-affiliated-proteins-rich profile for invasive ductal carcinoma to a more ECM glycoproteins/ECM regulators-rich profile for invasive lobular carcinoma (**Fig. 2B**).

In a third example, we used single-cell RNA-seq data obtained from 2,700 single peripheral blood mononuclear cells publicly available from 10X Genomics and used in the Seurat tutorial (Stuart et al., 2019). Pre-processing of the datasets identified nine clusters corresponding to the following cell types: B cells, memory CD4 T cells, naïve CD4 T, CD8 T cells, CD14^+^ monocytes, FCGR3A^+^ monocytes, NK cells, dendritic cells, and platelets. Tying the Seurat pipeline into Matrisome AnalyzeR enables the computation of the average expression of each gene for each single cell cluster, and the display as polar bar charts (or “`matristars`”) allows users to easily visualize the different matrisome categories arranged in a polar coordinate system, with the differences between categories being visualized through the length of their segments and height of their bars (**Fig. 2C**). Users can, at a glance appreciate the differential matrisome gene expression pattern across the different cell clusters, with B cells having the lowest number of ECM expressed genes (**Fig. 2C, left panel**) and platelets expressing a larger number of ECM genes encoding proteins involved in clotting (**Fig. 2C, right panel**). The complete analyzed dataset is available on the home page of the GitHub repository.

Upon completion of the workflow, users can extend their data analysis by using the output of the `matriannotate` and `matrianalyze` workflows to conduct comparative statistical analysis using the programs of their choice.

## DISCUSSION

The identification of genes or proteins belonging to the same functional compartment provides important information about the processes happening in cells and tissues and is a critical step in the analysis of large -omic datasets. Here, we report the deployment of a suite of versatile tools to annotate, classify, and tabulate ECM molecules in a variety of -omic datasets. Our goal was to develop tools accessible to non-ECM and ECM specialists alike, as well as novice and experts in big data analysis.

The current Matrisome AnalyzeR is designed to process data generated on the matrisomes of the five organisms the Naba laboratory and collaborators predicted. In recent years, others have predicted the avian (Huss et al., 2019), planarian (Cote et al., 2019; Sonpho et al., 2021), and bovine (Listrat et al., 2023) matrisomes. It is our goal to test the robustness of these predictions and evaluate their adoption by the scientific community. Should the number of -omic datasets on samples from these organisms increase, we will release augmented versions of Matrisome AnalyzeR to include these organisms as well.

Importantly, the field of “matrisomics” has significantly expanded in recent years, and we and others have developed additional tools to mine matrisomic datasets (Naba, 2023), such as MatrixDB, the database reporting ECM component interactions (http://matrixdb.univ-lyon1.fr/, (Berthollier et al., 2021; Clerc et al., 2019)), MatriNet, the database designed to explore network-scale changes in the ECM in pathophysiological conditions (https://www.matrinet.org/, (Kontio et al., 2022)), or the ECM proteomics database, MatrisomeDB (https://matrisomedb.org, (Shao et al., 2023)). It is our goal to deploy, in the future, releases of Matrisome AnalyzeR, that will create output that can directly be input to such databases to further advance ECM research and accelerate ECM biomarker discovery efforts.

## METHODS

### Matrisome lists of model organisms

The list of matrisome genes for the following model organisms were retrieved from their original publications: *Homo sapiens* (Naba et al., 2012a), *Mus musculus* (Naba et al., 2012a), *Danio rerio (Nauroy et al*., *2017), Drosophila melanogaster* (Davis et al., 2019), *Caenorhabditis elegans (Teuscher et al*., *2019)*. The lists are also available via the Matrisome Project website: https://sites.google.com/uic.edu/matrisome/matrisome-lists. The original gene identifiers were programmatically used to derive other general (NCBI gene, formerly Entrez Gene, and UniProt IDs) and species-specific identifiers: Ensembl Gene IDs for human and murine datasets, ZFIN IDs for zebrafish (Bradford et al., 2022), FlyBase ID for drosophila (Gramates et al., 2022), and WormBase and Common Gene Name for C. elegans datasets (Davis et al., 2022), using the annotation packages “org.Hs.eg.db”,”org.Mm.eg.db”, “org.Dr.eg.db”, “org.Ce.eg.db” and “org.Dm.eg.db”. The retrieved ID were finally manually reviewed and curated.

### Input file format stipulation

The only formatting requirement to files uploaded to the Matrisome AnalyzeR application is that they should contain column headers in their top row. Matrisome AnalyzeR accepts tab- and comma-separated (.tsv, .txt, .csv) files and is able to automatically recognize number format, though we encourage using dots for decimals and avoiding formatting thousands. The file size limit is 30 MB. If a file exceeds 30 MB, we recommend using the Matrisome AnalyzeR package.

If processing files using the Matrisome AnalyzeR package, the input format is a data.frame; the function will stop and issue a warning otherwise.

### Algorithms

The Matrisome AnalyzeR Shiny application and package are produced with the R Project for Statistical Computing and Shiny language (https://shiny.rstudio.com/) and share a common set of functions and “logic”. Users are expected to input a tabular dataset (typically, a high-throughput or - omic dataset) and identify a column with gene/protein identifiers and species information. Upon inputting the information, the first function of the pipeline (“`matriannotate`”) compares the input against a large database of matrisome annotations including gene symbols, NCBI gene (formerly Entrez Gene), and UniProt IDs for all species, as well as species-specific annotations such as Ensembl Gene IDs for human and murine datasets, ZFIN IDs for zebrafish datasets, FlyBase ID for *Drosophila* datasets, and WormBase and Common Gene Name for *C. elegans* datasets. Matching gene, protein or other ID are then enriched with matrisome divisions and categories (Naba et al., 2012a; Naba et al., 2012b), and non-matching values are returned as “non-matrisome”.

The output is organized to have the gene/protein/ID in the first column, followed by the annotated matrisome divisions, annotated matrisome categories, and the rest of the columns from the input file in their original order. This output is the base for the second function of the pipeline, “`matrianalyze`”, which takes in any numerical value in the dataset (also coercing resembling a number, *e*.*g*., values with “%” symbols, to a number) and sums them column-wise and by matrisome annotation. The result is a per-column (typically, per-sample) table of the quantity (*e*.*g*., number of reads, protein abundance, spectral counts) of any matrisome division and category across the entire dataset, which can be further used, for example, for statistical testing. The results from the “`matriannotate`” function are also the base for the graphical functions of both the application and package.

### Output file format

In the Matrisome AnalyzeR web application, the output on screen comprises a graphical and a tabular part. The graphical part is a bar chart, internally produced with the library ggplot2 (https://github.com/tidyverse/ggplot2) and customized to apply the color codes assigned to matrisome divisions and categories independently of the molecule IDs and species.

The tabular part is a browsable, scrollable, and searchable data table, internally produced with the library DT (https://rstudio.github.io/DT/). Upon completion of the “`matriannotate`” and/or “`matrianalyze`” functions, four download buttons appear in the navigation bar pointing to a single, zipped bundle including the tabular output in .csv format and the plot as a .pdf, or each of the outputs individually.

In the Matrisome AnalyzeR package, additional graphical functions are provided. These include donut charts (“`matrirings`”), polar bar chart (“`matristars`”), and Sankey/alluvial charts (“`matriflows`”). All graphs are internally produced with the library ggplot2 and with ggalluvial for `matriflows`. All graphical functions plot to screen by default, but this behavior can be changed by setting the “print.plot” parameter to FALSE. In this case, the underlying ggplot2 objects are returned instead, allowing further customization, integration with other pipelines, for example, printing to non-standard graphical devices. All tabular results are returned as data.frame.

## ACKNOWLEDGEMENTS

The authors would like to thank all the members of the Izzi and Naba laboratories for their feedback on Matrisome AnalyzeR and Monica Bassignana (www.monicabassignana.com) for her help with data visualization and the preparation of the graphs presented in the manuscript.

## DATA AVAILABILITY

All matrisome annotation lists are available at: https://sites.google.com/uic.edu/matrisome.

The web-based Matrisome AnalyzeR is deployed as a Shiny Application and is available at: https://sites.google.com/uic.edu/matrisome/tools/matrisome-analyzer.

The Matrisome AnalyzeR code package is available at: https://github.com/Matrisome/MatrisomeAnalyzeR.

## SUPPLEMENTARY MATERIAL LEGENDS

**Supplementary Table S1: Matrisome AnalyzeR_Test file**

.csv file containing proteomic data from three technical replicates on ECM samples from human Fallopian tubes (Renner et al., 2022) used as an example to demonstrate the functionalities of the web-based Matrisome AnalyzeR application.

**Supplementary Table S2:** .csv file resulting from inputting the test file provided as Supplementary File S1 and selecting the “Annotate” workflow of the web-based Matrisome AnalyzeR application.

**Supplementary Table S3_Analysis:** .csv file resulting from inputting the test file provided as Supplementary File S1 and selecting the “Annotate & Analyze” workflow of the web-based Matrisome AnalyzeR application.

**Supplementary Figure S1:** .pdf file containing bar graphs resulting from inputting the test file provided as Supplementary File S1 and selecting the “Annotate” workflow of the web-based Matrisome AnalyzeR application.

